# Hemispheric Complementarity is Associated with Distinct Functional and Structural Brain Profiles

**DOI:** 10.64898/2026.10.09.758042

**Authors:** Loïc Labache, Ahmad Beyh, Rachel Cooper, Linden Parkes, Avram J. Holmes

## Abstract

Hemispheric specialization is thought to follow a principle of complementarity, with homotopic functions such as language and attention lateralizing to opposite hemispheres, yet its consistency across individuals and relevance for cognition remain unclear. Using multimodal neuroimaging and twin-family data from 987 Human Connectome Project participants, we identified three sustained attentional lateralization phenotypes (right, left, bilateral) that were independent of handedness, differed by sex, and tracked broad shifts in brain organization, with bilateral individuals showing stronger intrahemispheric and homotopic connectivity. Examining attention and language revealed three multidomain profiles (crowded, mixed, split) reflecting whether functions colocalize within or are distributed across hemispheres. Mixed organization was linked to higher planum polare myelination, whereas crowded organization showed higher fiber density connectivity across dorsal attention–default network. Both phenotypes were heritable, yet neither predicted cognitive performance. These findings reveal a brain organization in which specialization emerges from flexible, heritable patterns linking function, structural connectivity, and myeloarchitecture.

---

Across species, the cerebral hemispheres exhibit systematic differences in their structure and function (1). These asymmetries constitute a fundamental organizational feature of the central nervous system, evident across molecular (2, 3), anatomical (4), and behavioral levels (5). The interdependence between a set of brain structures in one hemisphere and a given cognitive function is called hemispheric specialization (6). Yet, the principles underlying its coordination across functions remain unclear (7, 8). Language is typically left-lateralized (9–11), while attention; its homotopic counterparts (*i.e.*, anatomically corresponding across hemispheres (12)), involves a bilateral dorsal network principally contributing to goal-directed selection, and a right-lateralized ventral network mainly contributing to detect behaviorally relevant events and reorienting attention (13–16). The presence of hemispheric specialization is assumed to enhance efficiency by reducing redundancy and enabling parallel processing (6). The causal, or complementary, hypothesis proposes that lateralization of language to one hemisphere favors the lateralization of attention to the opposite hemisphere, thereby reducing interhemispheric competition (17). On the other hand, the statistical hypothesis proposes that their lateralization arises independently (7). However, evidence for their relationship is mixed: right-hemisphere language dominance has been associated with reversed attentional asymmetries (17, 18), but other studies report weaker, more bilateral attention responses (19) or partial dissociations between the two functions (20, 21). In this regard, how the hemispheric distributions of language and attention relate to one another, and more broadly how interindividual variability in their functional organization relates to cortical architecture, remains an open question.

A particularly informative window into this interindividual variability is provided by atypical hemispheric organization for language (*i.e.*, rightward dominance in lieu of leftward dominance), which is consistently observed in approximately 1 to 8% of the population (22–27). Atypical language lateralization is also associated with handedness, being disproportionately observed among left-handers, although the majority of left-handers nevertheless exhibit typical leftward language dominance (23, 24). Rather than simply being mirror-reversed copies of typical organization, atypical configurations exhibit distinct patterns of functional organization (22, 23). Instead, their language networks at rest show greater bilateral integration and stronger interhemispheric functional connectivity, while preserving a left-hemisphere network organization comparable to that of typical individuals, suggesting a left-language network organization invariant across typical and atypical individuals (23). More broadly, atypical language lateralization also appears to be reflected across multiple macroscale gradients and distributed associative systems, consistent with evidence that hemispheric specialization is organized along large-scale dimensions of cortical organization rather than isolated functional modules (22, 28–34). At the anatomical level, language lateralization has been associated with larger corpus callosum volume (23), and to intracortical myelin asymmetries of the auditory cortex (35, 36). Together, these findings emphasize the importance of moving beyond classical unimodal approaches to characterize hemispheric specialization through the integration of multiple brain features (6, 37).

Moving beyond language, individuals with stronger rightward lateralization of the ventral attentional system have been associated with stronger leftward language lateralization (38). However, the relationship is not straightforward and right-handed individuals typically exhibit complementary lateralization of language and attention, whereas left-handers show greater variability, including collateralized and co-dominant profiles (25). These observations raise the possibility that hemispheric specialization may emerge through multiple organizational processes rather than a single dominant developmental program (39). Sex also contributes modestly to variability in hemispheric organization, with population-level studies generally reporting slightly stronger structural and functional asymmetries in males than females, although effects are small and characterized by substantial overlap between sexes (23, 40). Beyond asymmetries observable through *in vivo* functional imaging, these organizational patterns may also be reflected in structural and microarchitectural features of the brain, including asymmetries in white matter pathways, cortical microstructure, and interhemispheric structural connectivity (28, 41, 42). Structural and functional asymmetries are moderately heritable and have been linked to genes involved in embryonic left-right axis formation, suggesting that variation in hemispheric specialization arises from shared neurodevelopmental mechanisms (22, 43, 44). Whether distinct patterns of language-attention lateralization correspond to separable neurobiological phenotypes differing in cortical microstructure, structural connectivity, and heritability remains to be established.

Here, we used multimodal neuroimaging and twin-family data from the Human Connectome Project (HCP) (45), to investigate how attention and language lateralization jointly define distinct profiles of hemispheric organization and how these profiles relate to functional and structural brain architecture. First, combining resting-state functional MRI (fMRI) with task activation studies of sustained attention, we identify three attentional lateralization phenotypes: typical right-dominant, bilateral, and atypical left-dominant. Individuals with bilateral attentional organization exhibit stronger within-network and homotopic functional coupling. Second, extending this framework across cognitive domains, we characterize the joint lateralization of attention and language through three organizational profiles: crowded, in which both functions lateralize to the same hemisphere; mixed, characterized by bilateral attentional organization; and split, in which the two functions lateralize to opposite hemispheres. Whole-brain analysis revealed that these profiles are associated with differences in cortical myeloarchitecture (left planum polare) and structural connectivity. Third, twin-based heritability analyses provide evidence that both attentional and crowding phenotypes are partly influenced by genetic factors. Together, these findings support the presence of multiple heritable, biologically grounded profiles of hemispheric organization, linking the lateralization of cognitive functions to variation in functional architecture, structural connectivity, and cortical microstructure.

## Results

### Population Variation in Attention Network Organization

To investigate hemispheric crowding, we first characterized individual variation in the lateralization of sustained attention. Crowding refers here to an organization in which cognitive functions that are typically dominant in opposite hemispheres instead become lateralized within the same hemisphere and recruit partly overlapping anatomical substrates, potentially increasing competition for shared neural resources (7, 31, 46).

First, attention lateralization was assessed in 987 participants of the Human Connectome Project cohort (45) with complete task and resting-state data (53% females, 11% left-handers). Individual measures were derived from the dorsal and ventral networks defined by the Atlas for Lateralized Attentional Networks (Fig. 1-b), or ALANs (14). Task-evoked asymmetry was calculated from the *2*-back *versus 0*-back contrast of the HCP working-memory task (47). Although the *N*-back paradigm primarily manipulates working-memory load, the HCP task-development study reported that the *N*-back elicited more robust activation of dorsal attention and cognitive-control regions than either event-related or blocked versions of a modified Posner attention task, both at the group level and across individual participants (47). We therefore do not interpret the *2*-back *versus 0*-back contrast as a measure of attentional orienting. Rather, we use increasing working-memory demand to probe the recruitment and hemispheric organization of the independently defined attention networks examined here. Accordingly, the *2*-back *versus 0*-back contrast provides a task-evoked measure of increasing goal-directed attentional-control demands (48, 49). Consistent with this interpretation, working-memory paradigms recruit attention-related networks (47), and *N*-back activation exhibits right-hemispheric biases within attention-related networks (50, 51). Goal-directed attention and working memory indeed share frontoparietal mechanisms (13). Experimental evidence shows that dorsal frontoparietal regions remain active while spatial information is maintained across a delay (52). We therefore used this contrast to characterize individual differences in the hemispheric organization of sustained, goal-directed attentional control under increasing cognitive demand. For each attentional network, asymmetry was quantified both across the complete network and within its principal hubs, regions that play a central role in network organization.

**Figure 1.**
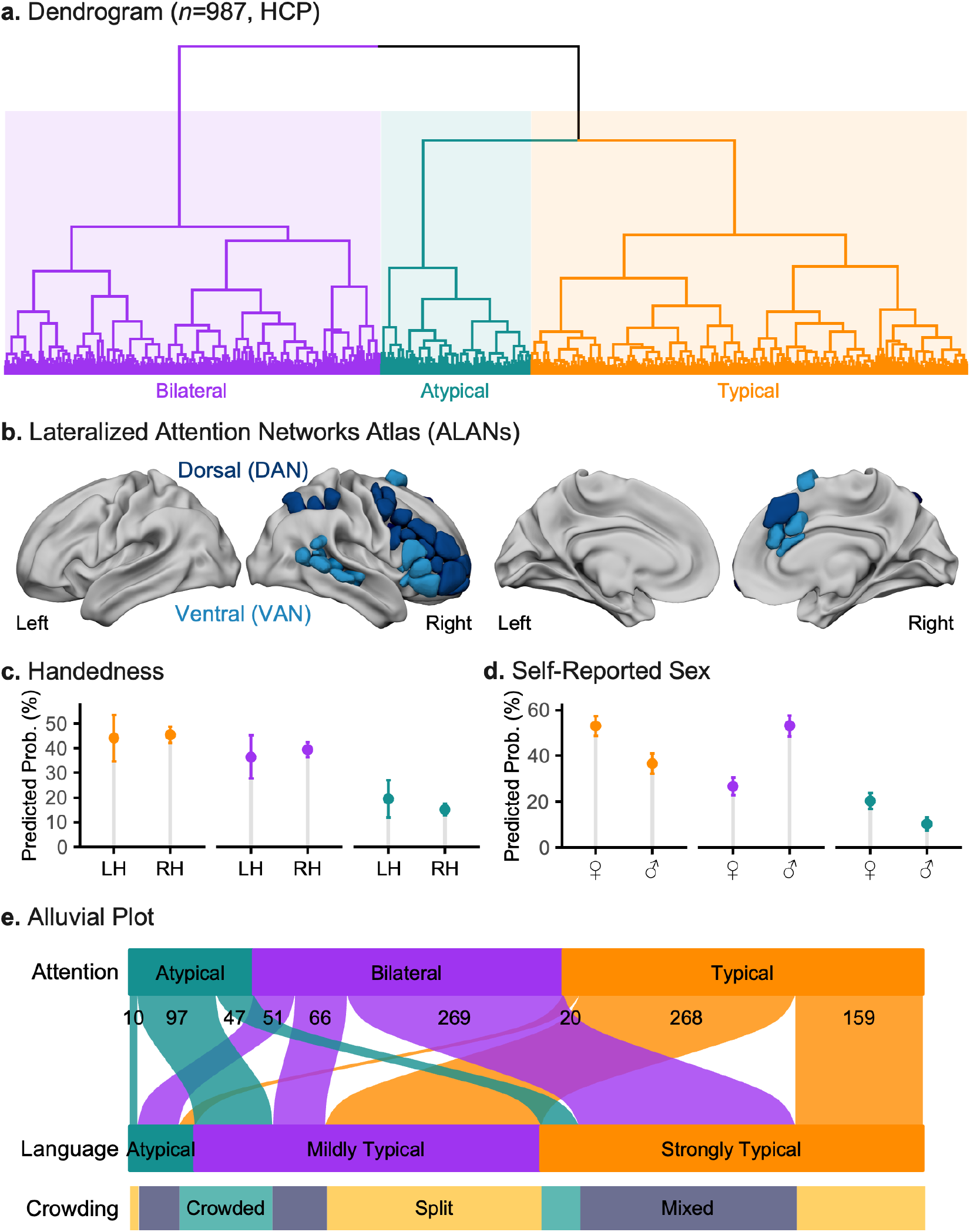
Population-level variation in attention and language lateralization reveals hemispheric crowding. **a,** Hierarchical clustering of individual-level lateralization indices in the Human Connectome Project sample (*n*=987) identified three attention phenotypes reflecting variation in hemispheric dominance. The Typical phenotype (orange; *n*=447, 46 left-handers, 278 females) showed rightward asymmetry, the Bilateral phenotype (purple; *n*=386, 42 left-handers, 140 females) showed no hemispheric dominance, and the Atypical phenotype (teal; *n*=154, 20 left-handers, 106 females) showed leftward asymmetry. Dendrogram branches are colored according to cluster membership. **b,** Lateralized Attention Networks Atlas (ALANs (14)) illustrating the two lateralized attention networks used to compute individual asymmetry measures: dorsal and ventral networks. Three-dimensional surface renderings of attentional regions are displayed on the HCP fs_LR (32k) template for the left and right hemispheres. **c-d,** Model-adjusted probabilities of attention phenotype membership by handedness (**c**) and self-reported sex (**d**). Predicted probabilities were obtained from an additive multinomial logistic regression model and averaged over the observed distribution of the other predictor. Self-reported sex was significantly associated with phenotype membership, likelihood-ratio *χ*^2^(2)=74.87, *p*<1.10^-3^. After adjustment for handedness, males were relatively more likely than females to exhibit bilateral rather than typical organization (*β*=1.06, SE=0.14, *p*<1.10^-3^; relative risk ratio RRR=2.88, CI_95%_=[2.17, 3.83]). Self-reported sex was not significantly associated with atypical rather than typical organization (*β*=-0.31, SE=0.20, *p*=0.13; RRR=0.74 ,CI_95%_=[0.50, 1.09]). Handedness was not significantly associated with phenotype membership (likelihood-ratio: *χ*^2^(2)=1.38, p=0.50). Error bars indicate 95% confidence intervals. LH, left-handed; RH, right-handed. **e,** Integration of attention phenotypes with previously defined language lateralization phenotypes (22) reveals three cross-domain configurations consistent with differing degrees of hemispheric complementarity: Split (*n*=437; 44 left-handers; 269 females), Mixed (*n*=386; 42 left-handers; 140 females), and Crowded (*n*=164; 22 left-handers; 115 females). Ribbon width reflects participant counts. Complementary organization (opposite hemispheric dominance across domains) predominates in the Split configuration, whereas collateralization characterizes the Crowded configuration.

The dorsal network hub was located in the inferior frontal sulcus, whereas the ventral network hubs encompassed the inferior frontal cortex, including the pars triangularis and anterior insula, and the posterior superior temporal sulcus (14). In the right hemisphere, the language network showed substantial spatial overlap with the combined attentional networks (Sørensen-Dice index (53, 54) SDI=0.49), indicatin hat the two cognitive systems share part of their anatomical substrate (14). This overlap was driven primarily by the ventral attention network (SDI=0.59), whereas overlap with the dorsal network was limited (SDI=0.12). Network-level asymmetries provide a marker with reduced interindividual variability, while hub-level asymmetries provide a more functionally specific measure of lateralization (9).

Task-derived measures were complemented by three resting-state measures for each network: mean intrahemispheric functional connectivity strength, asymmetry in functional connectivity strength, and homotopic interhemispheric connectivity. Thus, each participant was represented by ten measures describing the task-evoked and intrinsic functional organization of the two attentional networks. These data were entered into an agglomerative hierarchical clustering analysis following the procedures described previously (22, 23).

The unsupervised hierarchical clustering analysis supported a three-cluster solution, i.e. three attention phenotypes, with 12 of 21 *NbClust* (55) indices favoring three clusters (Fig. 1-a). The typical phenotype included 447 participants and was characterized by rightward task-evoked asymmetry (left-handers: *n*=46 (10%); female: *n*=278 (62%)). The bilateral phenotype included 386 participants and showed little evidence of consistent hemispheric dominance (left-handers: *n*=42 (11%); female: *n*=140 (36%)). The remaining 154 participants formed an atypical phenotype, characterized by leftward task-evoked asymmetry (left-handers: *n*=20 (13%); female: *n*=106 (69%)). The leftward attention profile accounted for 15.6% of this sample. These three multivariate profiles provide an operational description of variation in attention-network organization for the subsequent cross-domain analyses.

### Task-evoked asymmetry differentiates the attention phenotypes

Attention phenotype was associated with each of the four task-derived asymmetry measures after False Discovery Rate (FDR) correction (all *p*_FDR_<2.10^-27^, Fig. 2-b). Atypical participants showed leftward asymmetry in both attentional networks, at both spatial scales.

**Figure 2.**
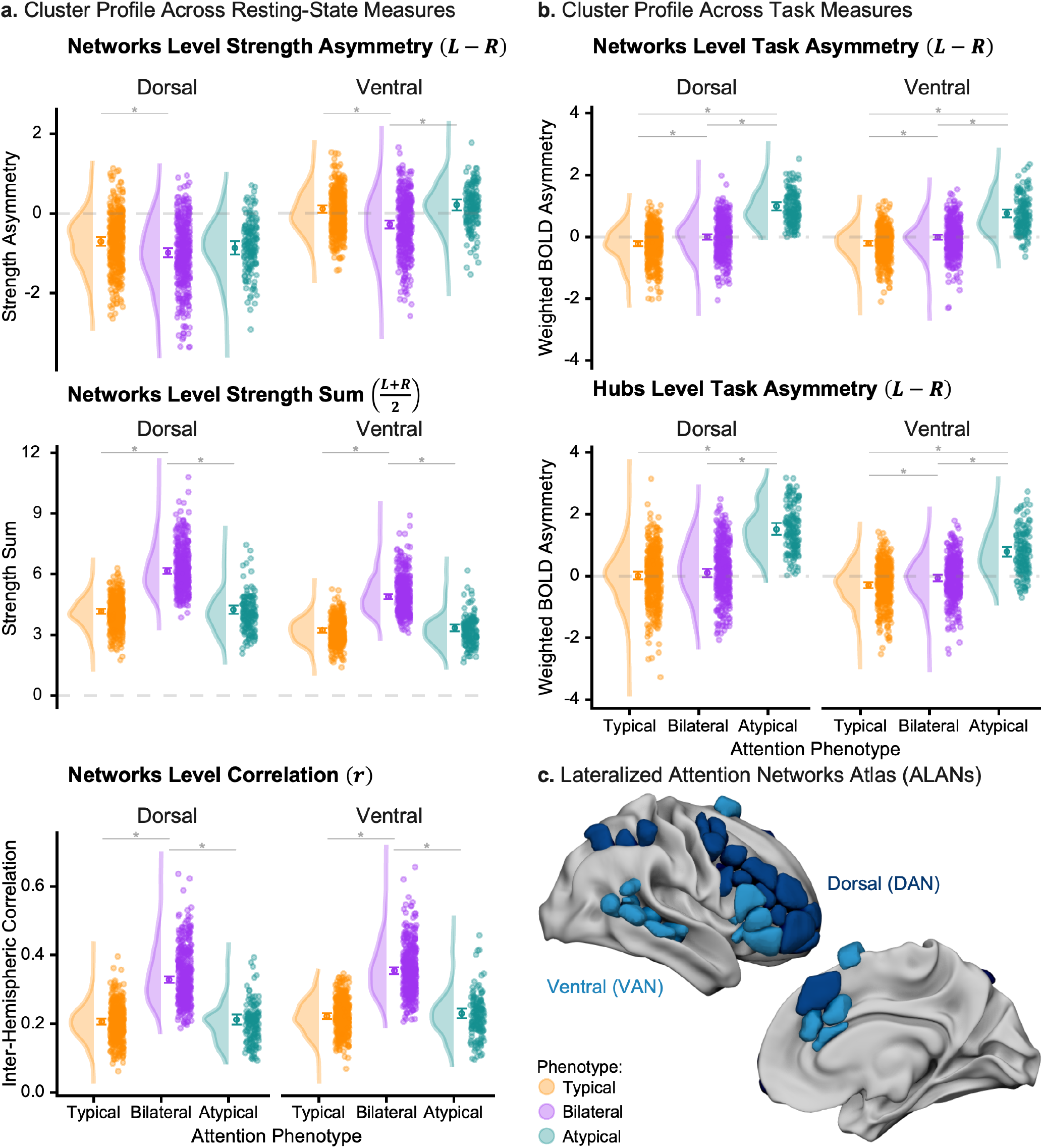
Multivariate hemispheric organization profiles underlying attention lateralization phenotypes. **a,** Resting-state network metrics across attention phenotypes. For dorsal and ventral networks, we quantified hemispheric strength asymmetry (*top*), bilateral network strength (*middle*), and interhemispheric functional connectivity (*bottom*). **b,** Task-evoked asymmetry measures for network-level and hub-level activation in the same networks. Across both resting-state and task measures, attention phenotype was significantly associated with hemispheric asymmetry (all *p*_FDR_<0.05). The Typical phenotype exhibited rightward asymmetry, the Atypical phenotype exhibited leftward asymmetry, and the Bilateral phenotype showed no hemispheric bias during task accompanied by increased interhemispheric connectivity and bilateral network strength. Points represent individual participants; solid circles indicate estimated marginal means ±95% confidence intervals from linear models. Asterisks denote effects surviving False Discovery Rate correction (*p*_FDR_<0.05). Positive asymmetry values indicate left hemisphere dominance. **c,** Lateralized Attention Networks Atlas (ALANs (14)) showing the dorsal and ventral networks, displayed on the HCP fs_LR (32k) template for the right hemisphere.

At the network level, the estimated marginal means were *μ*_network DAN_=0.99±0.13 for the dorsal network and *μ*_network VAN_=0.76±0.13 for the ventral network. At the hub level, the corresponding estimates were *μ*_hubs DAN_=1.52±0.20 and *μ*_hubs VAN_=0.78±0.16 , respectively. Here, μ denotes the estimated marginal means adjusted for the other predictors included in the model, and the values following denote the half-widths of the corresponding 95% confidence intervals.

*p*_FDR_ denotes the *p*-value for the main effect of attention phenotype using Benjamini-Hochberg false discovery rate correction.

In contrast, participants assigned to the typical phenotype showed rightward asymmetry at the network level in both the dorsal (*μ*_network DAN_=-0.22±0.09) and ventral (*μ*_network VAN_=-0.21±0.08) networks. Rightward asymmetry was also observed for the ventral network hubs (*μ*_hubs VAN_=-0.29±0.11). The dorsal network hub estimate was centered near zero (*μ*_hubs DAN_=0.01±0.13), indicating no clear hemispheric dominance at this spatial scale.

The bilateral group showed little task-evoked asymmetry overall. Network-level estimates were *μ*_network DAN_=-0.01±0.09 for the dorsal network and *μ*_network VAN_=-0.02±0.09 for the ventral network, while hub-level estimates were *μ*_hubs DAN_=0.10±0.14 and *μ*_hubs VAN_=- 0.06±0.11, respectively.

Atypical participants differed from both typical and bilateral participants across all four task-derived measures (all *p*_Tukey_<4.10^-13^). Typical participants also showed more rightward asymmetry than bilateral participants for three of the four measures (all *p*_Tukey_<7.10^-3^). The exception was the dorsal network hub measure, for which the typical and bilateral groups did not differ (*p*_Tukey_=0.61). Here, *p*_Tukey_ denotes the Tukey-adjusted post hoc pairwise comparisons between phenotype groups. Thus, the two lateralized profiles were not equal-magnitude mirror images. Weak task asymmetry also did not uniquely identify the Bilateral phenotype, whose distinction became clearer when its intrinsic connectivity profile was considered.

### Intrinsic connectivity distinguishes bilateral attention organization

The analyses above described how the phenotypes differed in evoked asymmetry. We next characterized the resting-state features that are also linked to the same clustering solution. This complements prior evidence that language lateralization is associated with distributed intrinsic connectivity organization (22), while asking how intrinsic network strength and interhemispheric coupling distinguish the attention profiles defined here.

Attention phenotype was associated with connectivity-strength asymmetry in both the dorsal and ventral networks (all *p*_FDR_<4.10^-3^, Fig. 2-a). The clearest separation was observed in the ventral network. Bilateral participants showed rightward connectivity-strength asymmetry (*μ*_strength asym VAN_=-0.27±0.10), different from both typical and atypical participants (both *p*_Tukey_<1.10^-7^). Typical (*μ*_strength asym VAN_=0.11±0.09) and atypical (*μ*_strength asym VAN_=0.21±0.15) participants instead showed modest leftward asymmetry and did not differ from one another (pTukey=0.44).

For the dorsal network, bilateral participants showed greater rightward connectivity-strength asymmetry (*μ*_strength asym DAN_=-0.98±0.12) than typical participants (*μ*_strength asym DAN_=-0.70±0.11, *p*_Tukey_=2.10^-3^). The estimate for atypical participants was intermediate (*μ*_strength asym DAN_=-0.86±0.17), and comparisons involving the atypical group were not significant (all *p*_Tukey_>0.27).

Attention phenotype was also strongly associated with total connectivity strength and homotopic interhemispheric connectivity in both attentional networks (all *p*_FDR_<5.10^-65^, Fig. 2-a). For these measures, typical and atypical participants exhibited comparable profiles (all *p*_Tukey_>0.52). In the dorsal network, typical participants had a total connectivity-strength estimate of *μ*_strength sum DAN_=4.46±0.13 and an interhemispheric-connectivity estimate of *μ*_r DAN_=0.21±0.01, compared with *μ*_strength sum DAN_=4.24±0.21 and *μ*_r DAN_=0.21±0.02, respectively, among atypical participants. In the ventral network, the corresponding estimates were *μ*_strength sum VAN_=3.23±0.11 and *μ*_r VAN_=0.22±0.01 for typical participants and *μ*_strength sum VAN_=3.34±0.17 and *μ*_r VAN_=0.23±0.01 for atypical participants.

The bilateral phenotype was distinguished by substantially higher values on both measures. Bilateral participants had total-strength and interhemispheric-connectivity estimates of *μ*_strength sum DAN_=6.16±0.15, and *μ*_r DAN_=0.33±0.01, respectively, in the dorsal network, and *μ*_strength sum VAN_=4.48±0.12 and *μ*_r VAN_=0.35±0.01 in the ventral network. Each of these values was higher than the corresponding estimates in both the typical and atypical groups (all *p*_Tukey_<4.10^-13^). Thus, the bilateral phenotype was not simply intermediate between the two lateralized phenotypes. Instead, it was characterized by stronger within-network connectivity and greater coupling between homotopic regions across hemispheres.

### Attention Phenotype Associations with Self-Reported Sex and Handedness

Because self-reported sex and handedness have both been implicated in interindividual variation in hemispheric specialization, we next asked whether they were associated with attention phenotype membership. Prior work has identified small population-level sex differences in functional cerebral asymmetries, including visuospatial lateralization, whereas handedness has been linked to variation in spatial-attentional biases and atypical patterns of hemispheric organization (8, 23, 56). We therefore examined whether the attention phenotypes were related to these previously implicated sources of variation in lateralization. Adding the self-reported sex by handedness interaction did not improve model fit, likelihood-ratio *χ*^2^(2)=0.23, *p*=0.89. We therefore examined the main effects of self-reported sex and handedness using an additive multinomial logistic regression model. Self-reported sex was significantly associated with attention phenotype membership (Fig. 1-d), likelihood-ratio *χ*^2^(2)=74.87, *p*<2.10^-16^. After adjustment for handedness, males were relatively more likely than females to exhibit bilateral rather than typical organization (*β*=1.06, SE=0.14,

*p*=2.10^-13^; relative risk ratio RRR=2.88, CI_95%_=[2.17, 3.83]). Self-reported sex was not significantly associated with membership in the atypical rather than the typical phenotype (*β*=-0.31, SE=0.20, *p*=0.13; RRR=0.74, CI_95%_=[0.50, 1.09]). Handedness was not significantly associated with attention phenotype membership (Fig. 1-c), likelihood-ratio *χ*^2^(2)=1.38, *p*=0.50. The sex association primarily concerned bilateral rather than rightward attention organization, whereas neither the leftward versus rightward contrast by sex nor the overall handedness effect was detected. Thus, the demographic pattern differed from the previously reported enrichment of atypical language organization among left-handers in this cohort (22) and in an independent cohort (23).

### Population Variation in Hemispheric Complementarity

Having characterized individual variation in attention lateralization, we next examined whether the hemispheric organization of attention and language followed a coordinated pattern consistent with complementary specialization. The causal or complementary specialization hypothesis predicts that lateralization of one function should favor allocation of the other function to the opposite hemisphere, thereby reducing potential competition within shared cortical systems (17). By contrast, the statistical hypothesis predicts that the lateralization of different functions should arise relatively independently (7).

To relate the newly derived attention phenotypes to language organization, we used the previously published language assignments for the same HCP participants (22). That analysis combined two story versus math activation asymmetries with three intrinsic connectivity measures in a separate clustering of the language network. It identified strong typical language, with pronounced leftward asymmetry; mild typical language, with more moderate leftward asymmetry; and atypical language, with rightward asymmetry. Restricting those existing assignments to the 987 participants studied here yielded 475 strong typical, 431 mild typical and 81 atypical language profiles (Fig. 1-e).

We next mapped the two domain-specific classifications (language and attention) onto three cross-domain configurations without reclustering participants. For this mapping, strong typical and mild typical language both indicated left-dominant language, whereas atypical language indicated right-dominant language. Split denotes opposite dominant hemispheres for attention and language, and Crowded denotes the same dominant hemisphere. Mixed denotes bilateral attention regardless of language phenotype. The resulting groups comprised 437 Split participants (44.3%, 44 left-handers, 269 females), 386 Mixed participants (39.1%, 42 left-handers; 140 females), and 164 Crowded participants (16.6%, 22 left-handers, 115 females). These categories correspond to the cross-domain configurations illustrated in Fig. 1-e.

At the full three-level resolution, attention and language phenotype assignments were significantly associated (Pearson’s χ2(4, n=987)=184.50, p<1.10-15, Cramér’s V ϕC=0.31). Analyses of the nine attention-language phenotype combinations showed that the Typical attention phenotype was enriched among individuals with Mild Typical language organization (adjusted standardized residual, *Adj. res*=9.39) and depleted among individuals with Strong Typical (*Adj. res*=-7.18) or Atypical language organization (*Adj. res*=-3.89). The Bilateral attention phenotype showed the inverse pattern, with enrichment among individuals with Strong Typical (*Adj. res*=10.87) and Atypical language organization (*Adj. res*=4.59), and depletion among those with Mild Typical language organization (*Adj. res*=-13.49). Finally, the Atypical attention phenotype was enriched among individuals with Mild Typical language organization (*Adj. res*=5.26) and depleted among those with Strong Typical language organization (*Adj. res*=-4.76). All of these effects remained significant after FDR correction (all pFDR<1.10-4). The number of participants jointly exhibiting Atypical attention and Atypical language organization did not differ from that expected under independence (*Adj. res*=-0.84, pFDR=0.40).

The three by three analysis tested association between the full multivariate phenotypes. We then asked a narrower question: among participants assigned a lateralized attention phenotype, were the dominant hemispheres for language and attention associated? For this two by two analysis only, we excluded the bilateral attention group and pooled strong typical and mild typical language as left-dominant, retaining atypical language as right-dominant. This recoding used the same domain-specific assignments and did not introduce a new clustering solution. Among the remaining 601 participants, 437 (72.71%) exhibited complementary, opposite-hemisphere organization and were classified as Split, whereas 164 (27.29%) exhibited same hemisphere organization and were classified as Crowded. Despite this apparent predominance of Split organization, language dominance was not significantly associated with attention dominance after accounting for the marginal frequencies of the phenotypes (χ2(1, n=601)=0.98, p=0.32, ϕC=0.04). The point estimate suggested higher odds of complementary split organization, but the confidence interval was wide and included both no association and effects in the opposite direction (odds ratio OR=1.48, CI95%=[0.68, 3.24]), with values above 1 indicating greater odds of rightward attention in left-language than in right-language participants.

The full multivariate language and attention classifications were associated, whereas the directional test did not detect an association between their dominant hemispheres. The high prevalence of Split organization therefore does not itself demonstrate coupling beyond the marginal frequencies of leftward language and rightward attention. The cross-classification instead highlights an association involving bilaterality and the distinction between strong typical and mild typical language. However, the directional confidence interval was broad, and this result does not establish directional independence or exclude probabilistic complementarity. These findings are inconsistent with both a causal deterministic complementarity hypothesis and statistical independence, and instead suggest the presence of multiple population-level patterns of hemispheric organization.

### Heritability of Attention and Crowding Phenotype

To determine whether these population-level phenotypes have a genetic component, we estimated the heritability of attention lateralization and cross-domain crowding using the family structure of the HCP cohort. Heritability was estimated using the multidimensional heritability analysis approach as described by Anderson and colleagues (57). The attention-lateralization phenotype showed an estimated heritability of *h*^2^=12.51% (SE=4.94%, *p*=0.003). Follow-up analyses indicated that crowding phenotype was also significantly heritable (*h*^2^=13.22%, SE=4.93%, *p*=0.003). This suggests that both the lateralization of the attention networks and hemispheric complementarity are under genetic control. For reference, language lateralization in this cohort was previously reported to be heritable at *h*^2^=11.2% (22). Collectively, these results suggest that neuronal asymmetries likely developed under phylogenetic pressure, and therefore possess a genetic basis (58).

### Comparable Cognitive Performance Regardless of Crowding

To determine whether differences in hemispheric organization were accompanied by measurable differences in cognitive performance, we compared broad and domain-specific cognitive measures across the crowding phenotypes. Analyses included 974 participants with complete phenotype, cognitive, and covariate data. After adjustment for manual preference, age, years of education, total intracranial volume, and self-reported sex, crowding phenotype was not significantly associated with the Total Cognition Composite (*p*_FDR_=0.65, *η*_P_^2^=3.10^-3^), the Crystallized Cognition Composite (*p*_FDR_ =0.67, *η*_P_ ^2^=2.10^-3^), or the Fluid Cognition Composite(*p*_FDR_=0.73, *η*_P_^2^=1.10^-3^). No differences among the Split, Mixed, and Crowded phenotypes were observed for visual episodic memory, cognitive flexibility, inhibitory control, reading decoding, vocabulary comprehension, processing speed, or working memory (all *p*_FDR_≥0.23, all *η*_p_ ^2^≤3.10^-3^). Finally, the association between crowding phenotype and cognitive performance was neither modulated by manual preference for any composite or individual cognitive measure (all *p*_FDR_>0.87, all *η*_p_ ^2^≤3.10^-3^). Thus, despite significant differences in the hemispheric allocation of language and attention, the Split, Mixed, and Crowded phenotypes showed no detectable differences in broad or domain-specific cognitive performance.

### Mixed Organization is Associated with Higher Planum Polare Cortical Myelination

Given prior evidence linking language lateralization to intracortical myelin (35), we next examined whether the crowding phenotype was associated with local variation in cortical myeloarchitecture. At each cortical vertex, myelin-map values were residualized for local cortical thickness and curvature. The resulting residuals were entered into a whole-cortex vertex-wise analysis controlling for age, self-reported sex, total intracranial volume, and manual preference. Random-field-theory-corrected analysis identified a single positive cluster in the left anterior superior temporal and para-insular cortex, centered on the planum polare (179 vertices, *p*_RFT_=1.10^-3^, *t*^peak^=5.89, MNI coordinates: *x*=-45.7, *y*=-9.4, *z*=-6.6, Fig. 3-a). The planum polare occupies the anterior supratemporal plane, anterior to Heschl’s gyrus and the planum temporale. In the 17 networks parcellation proposed by Yan and colleagues (59), the planum polare is assigned to the auditory network. Mean residualized myelin values extracted from this cluster instead differed across the three crowding phenotypes (*p*=1.10^-8^, Fig. 3-b). Tukey-adjusted comparisons indicated that the Mixed phenotype (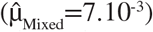) had a significantly higher (both *p*<1.10^-4^) adjusted myelin values than both the Split phenotype (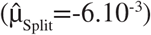) and the Crowded phenotype (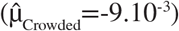). Split and Crowded participants did not differ significantly (*p*=0.54). Thus, local myelin variation was not monotonic with the degree of hemispheric crowding. Instead, increased myelin map values within the left planum polare were selectively associated with Mixed organization, characterized by bilateral sustained-attention organization.

**Figure 3.**
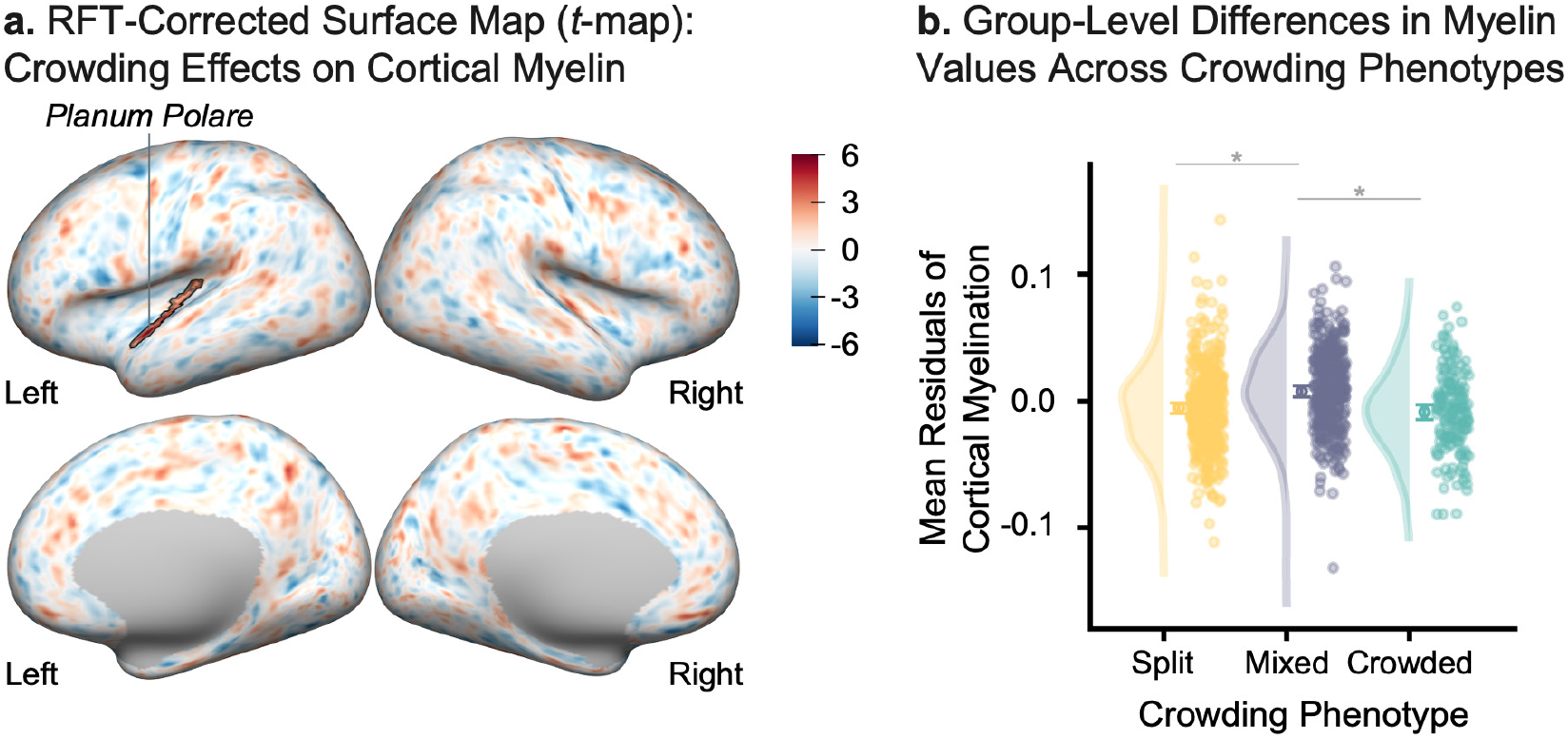
Hemispheric crowding is associated with focal differences in cortical myelination. **a,** Surface-based vertex-wise analysis of residualized cortical myelin values as a function of crowding phenotype. Random field theory correction identified one positive significant cluster in the left anterior superior temporal and para-insular cortex, centered in the planum polare (179 vertices, *p*_RFT_=1.10^-3^, *t*_peak_=5.89, MNI coordinates: *x*=-45.7, *y*=-9.4, *z*=-6.6). Black outlines indicate vertices belonging to the significant cluster. **b,** Mean residualized myelin values extracted from the significant cluster across crowding phenotypes (Split, Mixed, Crowded). Points represent individual participants; solid circles denote estimated marginal means ±95% confidence intervals from linear models. The Mixed phenotype had higher values than the Split and Crowded phenotypes (both *p*<1.10^-4^). Split and Crowded participants did not differ (*p*=0.54). Pairwise comparisons were adjusted using Tukey’s range test.

### Crowded Organization is Associated with Higher Fiber Density across Distributed Connections

Hemispheric specialization is supported not only by the functional segregation of cortical systems but also by the white-matter pathways that coordinate processing within and between hemispheres (28, 60, 61). We therefore asked whether the cross-domain configuration of language and attention was accompanied by differences in structural connectivity. Structural connectivity was quantified using median hindrance modulated orientational anisotropy (HMOA (62)), used here as an index of fiber density, between 400 cortical parcels assigned to seven canonical functional networks (Fig. 4-a). Of the 79,800 possible undirected parcel-to-parcel connections, 62,182 had nonzero HMOA values and were included in the edge-wise analyses; 385 showed a significant main effect of crowding phenotype after FDR correction (all *p*_FDR_<0.05, Fig. 4). These connections were distributed across 26 of the 28 possible within- and between-network combinations, indicating that differences associated with hemispheric crowding were broadly distributed rather than restricted to a single anatomical pathway or functional system.

**Figure 4.**
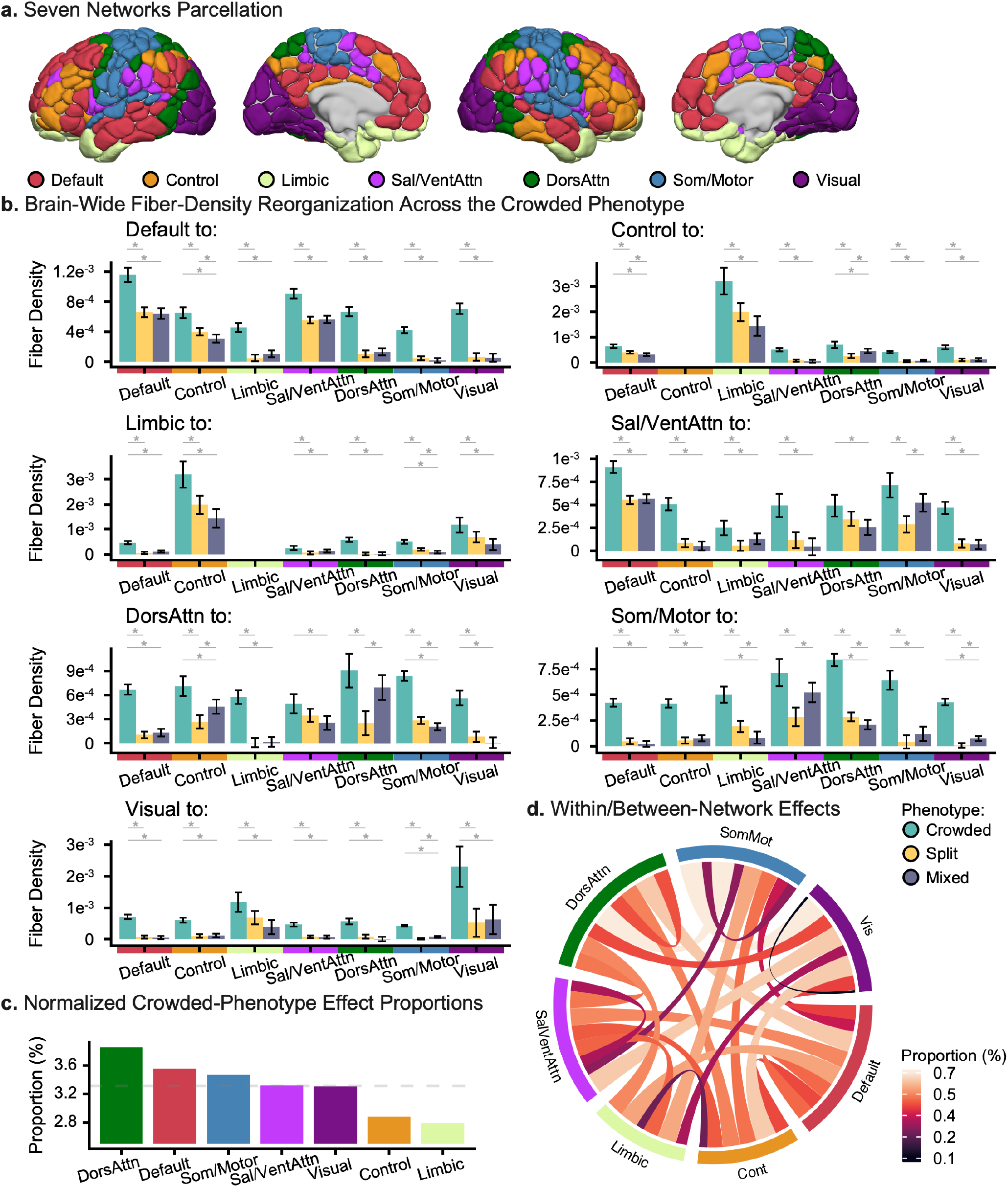
Crowding phenotype is associated with distributed increases in fiber density across large-scale networks. **a,** Seven network cortical parcellation (63) mapped onto 400 homotopic parcels (59) used for structural connectivity analyses. **b,** Network-wise least-squares mean fiber density across crowding phenotypes (Crowded, Split, Mixed). For each source network (“*Network* to”), bars represent adjusted marginal means (±95% CI) of fiber density projected to all target networks, estimated from linear models. Asterisks denote significant pairwise differences surviving False Discovery Rate correction (*p*_FDR_<0.05). Across multiple source networks, the Crowded phenotype exhibited higher fiber density relative to Split and Mixed phenotypes. **c,** Normalized proportion of edges showing a significant Crowded phenotype effect by network. Network involvement was summarized by computing, for each network pair, the proportion of possible connections showing a significant crowding effect and then summing these pairwise normalized edge densities across all pairs involving each network. Dorsal Attention and Default networks contributed the largest proportion of altered connections. **d,** Chord diagram summarizing within- and between-network effects. Ribbon thickness and color encode the proportion of significant edges associated with the Crowded phenotype. Effects were distributed across both intra- and inter-network connections, indicating widespread alterations in structural connectivity.

To characterize the direction of these effects, we averaged HMOA values across the significant edges belonging to each undirected network pair and calculated covariate-adjusted estimated marginal means for the three crowding phenotypes. The Crowded phenotype exhibited higher fiber density than the Split phenotype in 25 of the 26 network pairs (96.2%) and higher fiber density than the Mixed phenotype in 24 of the 26 pairs (92.3%; *p*_Tukey_<0.05, Fig. 4-b). No network pair showed significantly lower fiber density in Crowded participants. By contrast, Split and Mixed participants differed in only 7 of the 26 network pairs: fiber density was higher in Split participants for three pairs, higher in Mixed participants for four pairs, and did not differ significantly for the remaining 19 pairs (73.1%). The structural-connectivity effect was therefore not characterized by a monotonic progression from Split to Mixed to Crowded organization. Instead, it reflected a distributed elevation in fiber density that was specifically associated with the Crowded phenotype.

After accounting for differences in network size and in the number of possible parcel-to-parcel connections, the dorsal attention network showed the greatest normalized involvement, followed by the default and somatomotor networks (Fig. 4-c). Prominent between-network effects included connections linking the dorsal attention and somatomotor networks, the somatomotor and visual networks, and the default and dorsal attention networks. Significant effects encompassed both within- and between-network connections (Fig. 4-d). Together, these findings indicate that collateralization of language and attention within the same hemisphere is associated with widespread increases in HMOA-weighted structural connectivity, with particularly strong involvement of attention-related and trans-modal systems.

## Discussion

Hemispheric specialization is commonly described in terms of population-level biases, most prominently leftward language and rightward attention. Yet these canonical patterns obscure substantial interindividual variation and leave unresolved whether lateralized cognitive functions are organized independently or according to coordinated principles of hemispheric allocation. Here, we identified three phenotypes of sustained-attention organization and showed that their relationship with language lateralization could not be reduced either to strict complementary specialization or to complete statistical independence. Instead, language and attention were coupled primarily through differences in the strength and bilaterality of hemispheric organization. These cross-domain configurations were heritable and associated with distinct cortical microarchitectural and white-matter profiles, yet were not accompanied by detectable differences in broad cognitive performance. Together, these findings suggest that hemispheric specialization may be better understood as a set of biologically grounded population-level configurations than as variation around a single canonical architecture.

Variation in attention lateralization was organized into right-dominant, bilateral and left-dominant phenotypes. Importantly, Bilateral participants were not simply intermediate between the two lateralized groups. This group showed little consistent hemispheric bias during task performance, but substantially greater within-network connectivity and homotopic interhemispheric coupling at rest. This dissociation indicates that reduced task lateralization can coexist with a highly integrated intrinsic architecture. Task-evoked asymmetry and resting-state connectivity should therefore not be treated as interchangeable measures of hemispheric dominance. Consistent with this distinction, the canonical organization of attention systems is evident in spontaneous activity, with intrinsic correlations revealing a largely bilateral dorsal system alongside a right-lateralized ventral system (64). Previous findings in language similarly indicate that task-evoked dominance need not map directly onto intrinsic network organization, with atypical individuals showing no dominant hemisphere at rest (23). Furthermore, bilateral organization at rest can reflect homotopic coupling and intrinsic network architecture even when task engagement produces clearer hemispheric specialization (65). More broadly, association cortex contains both specialized and functionally flexible territories, with flexible regions positioned to integrate information across specialized functions (66). The bilateral phenotype may therefore reflect a distinct integrative organization rather than merely weakened rightward specialization. This interpretation is consistent with broader accounts of hemispheric connectivity. Forexample, previousworksuggeststhatright-lateralized processes tend to participate in comparatively distributed and interhemispheric interactions, whereas left-lateralized functions show stronger intrahemispheric organization (67). At the same time, lateralization does not appear to reflect a unitary tendency for an individual to be globally left- or right-dominant. Rather, asymmetries are organized across partially independent functions (32, 68). The present findings extend this framework by demonstrating substantial population variation within attention-related networks themselves, including a common bilateral configuration characterized by increased intrinsic integration.

The demographic associations further argue against reducing attentional lateralization to other familiar markers of hemispheric specialization. Handedness was not associated with attention phenotype, consistent with evidence that manual preference has a complex and limited relationship with visuospatial lateralization. Although strong left-handers can show distinctive relationships between language and attention asymmetry, handedness does not generally determine the lateralization of visuospatial functions (8, 69). Self-reported sex, by contrast, was associated with attention phenotype, with males more likely than females to exhibit bilateral rather than typical organization. This effect should be interpreted as a population-level association rather than evidence for discrete sex-specific forms of hemispheric organization, given the substantial within-group variability reported for structural and functional brain asymmetries (40).

The relationship between language and attention provides a more direct test of competing accounts of complementary specialization and statistical independence (17, 70). At the full phenotypic level, their organization was clearly non-independent. Strong typical language organization was preferentially associated with bilateral attention, whereas mild typical language organization was enriched among the lateralized attention phenotypes. However, when the analysis was restricted to individuals with an identifiable attention-dominant hemisphere, language dominance did not predict the direction of attention dominance. Thus, the association between the two functions was expressed primarily through lateralization strength and bilaterality, rather than through a deterministic rule whereby assigning one function to a hemisphere forces the other into the contralateral hemisphere. This result helps reconcile apparently conflicting findings in the literature. Cai and colleagues (17) reported complementary language-attention lateralization in a sample enriched for typical and atypical language dominance, whereas other studies found weaker or population-dependent associations. Zago and colleagues (71), for example, observed a negative association between language and attention lateralization specifically among strong left-handers, while both empirical studies and reviews indicate substantial heterogeneity in the joint lateralization of language and visuospatial functions, including complementary, reversed, bilateral, and crowded configurations (72, 73). Taken together, these findings suggest that language and attention lateralization are neither fully independent nor always complementary. Their relationship appears to depend more on the strength and bilaterality of hemispheric organization than on a fixed rule assigning the two functions to opposite hemispheres. Complementary organization may therefore represent a common population-level configuration rather than a deterministic relationship between the hemispheric dominance of the two functions. The prevalence of different organizational motifs reinforces this interpretation. Among participants with lateralized attention, split organization was more frequent than crowded organization, yet more than one quarter showed same-hemisphere lateralization of language and attention. Across the full sample, a further large fraction exhibited bilateral attention and was classified as mixed. Earlier estimates of crowding have varied widely according to task, handedness, sample composition and operational definition, with healthy cohorts showing both complementary and non-complementary organizations (71, 72, 74). In the present large HCP cohort, alternative cross-domain configurations were sufficiently frequent that they are difficult to regard simply as rare deviations from a canonical phenotype. Future studies should incorporate multiple language and attention paradigms within the same samples to establish whether these cross-domain configurations are robust across task contexts and different definitions of hemispheric specialization.

The heritability of both attention phenotype and cross-domain crowding further indicates that this population variation has a developmental biological component. The magnitude of these estimates was similar to the previously reported heritability of language lateralization in the same cohort (22). Contemporary accounts instead suggest that genetic influences bias developmental trajectories while structural maturation, experience and functional plasticity continue to shape the resulting architecture (65). This is particularly relevant because different functional asymmetries mature along distinct trajectories. Language networks progressively transition from strong interhemispheric coordination toward increasing left intrahemispheric organization, while rightward frontoparietal asymmetry also develops across childhood (8). The adult phenotypes identified here may therefore represent alternative developmental trajectories rather than fixed states established early in life.

The cortical myelin findings provide evidence that these organizational differences are accompanied by local microarchitectural variation. The only significant cortical effect was centered on the left planum polare, where mixed participants showed higher T1w/T2w-derived myelin values than either split or crowded individuals. Split and crowded participants did not differ, arguing against a monotonic relationship between crowding and cortical myelination. Instead, the effect was specifically associated with the configuration characterized by bilateral attention. The localization to anterior superior temporal cortex is notable given the role of superior temporal regions in auditory and language-related processing (9, 75), but its functional interpretation should remain cautious. Recent work further links cortical myeloarchitecture to hierarchical differences between phonological and semantic language processing (76). The present result therefore links cross-domain lateralization to cortical microstructure, but does not establish whether increased T1w/T2w values represent a developmental antecedent, adaptation, or other tissue property associated with bilateral organization. This caution is reinforced by the lifespan dynamics of cortical myelination. Myelin-sensitive MRI measures vary markedly across cortical regions and follow nonlinear developmental and aging trajectories (77). The restricted age range of the HCP cohort therefore provides only a cross-sectional view of a potentially drawn out developmental process. Determining when the planum polare difference emerges will be important for distinguishing whether it precedes, accompanies or follows the establishment of bilateral sustained attention. This question is especially relevant given evidence that functional and connectivity-based asymmetries can mature on different timescales (8) and the developmental and environmental contributors to hemispheric specialization remain to be fully characterized. A distributed anatomical signature emerged in white matter. Crowded participants showed higher HMOA-weighted structural connectivity across nearly every network pair exhibiting a significant phenotype effect, with particularly strong normalized involvement of dorsal attention, default and somatomotor networks. This pattern argues against crowding being supported by alteration of a single canonical tract. Instead, same-hemisphere organization of language and attention appears embedded in a broader structural-connectivity architecture. Earlier work reported increased callosal anisotropy in individuals with atypical hemispheric segregation, particularly when language and spatial attention were crowded into the same hemisphere (78). More broadly, white-matter pathways implicated in language and visuospatial attention show substantial asymmetry and interindividual variability, while anatomical lateralization does not necessarily provide a straightforward marker for functional lateralization (4, 79, 80). One possibility is that increased structural connectivity provides additional routes for coordinating cognitive functions that occupy overlapping hemispheric territories. Such an interpretation would be compatible with the idea that long-range structural architecture provides a relatively persistent anatomical scaffold that constrains functional lateralization while remaining itself plastic across development and experience (65). It would also fit with the increased intrinsic integration observed in bilateral participants.

Despite these neurobiological differences, cross-domain organization was unrelated to broad cognitive performance. This null result is informative because hemispheric specialization is often proposed to enhance processing efficiency by separating computations across hemispheres and reducing competition between neural resources (46, 81). The empirical evidence for a generalized cognitive advantage of canonical lateralization is nevertheless mixed (72). Previous studies have reported associations between the strength of lateralization and cognitive performance (81, 82), whereas others have found no detectable behavioral disadvantage associated with atypical or crowded configurations (81, 83–85). Developmental work has similarly found that crowding language and visuospatial functions within the same hemisphere was not associated with poorer cognitive or academic performance (83–85). The present results extend this evidence to a large adult cohort across multiple NIH Toolbox domains. The absence of broad cognitive differences does not imply that hemispheric organization is independent from behavior. One possibility is that the costs of crowding emerge primarily when lateralized functions must operate concurrently and compete for overlapping neural resources, rather than during standard tests that assess cognitive domains separately. Consistent with this possibility, Zhu and Cai (86) found that individuals with co-lateralized language and spatial attention showed poorer overall performance and greater interference than individuals with the typical complementary pattern during a dual-task combining verbal fluency and spatial processing, whereas these group differences were not evident in the corresponding single-task conditions (86). This distinction is important because the NIH Toolbox indexes broad abilities largely under isolated task conditions. More generally, prior work suggests that the magnitude of lateralization may sometimes be more closely related to performance than its direction, and that relationships between lateralization and cognition depend strongly on the function, population, and measurement approach (72, 81). Thus, the phenotypes identified here may differ in processing strategy or susceptibility to concurrent interference without producing detectable differences in overall cognitive ability. Equivalent behavioral performance across markedly different neural configurations is also compatible with the principle of degeneracy, whereby distinct neural processes can support similar functions (87, 88). This provides a useful alternative to interpreting bilateral, atypical or crowded organization as intrinsically deficient. The increased structural connectivity of crowded participants and increased intrinsic integration of bilateral participants may represent different architectures capable of preserving similar behavioral output.

Several limitations are important. First, the task-derived attention measures were based on the 2-back versus 0-back HCP working-memory contrast. Although this contrast recruits the independently defined attention networks examined here, it manipulates working-memory load and goal-directed cognitive control rather than providing a process-pure measure of visuospatial attention (48, 49). The task component of these phenotypes should therefore be interpreted as reflecting the hemispheric organization of sustained, goal-directed attentional control under increasing cognitive demand. Hemispheric asymmetries themselves vary with task, modality and difficulty (69), and future work should test whether the same phenotypes emerge during spatial orienting or target-detection paradigms. Finally, the HCP cohort comprises primarily healthy young adults, limiting inference about the developmental origin, aging trajectory and clinical relevance of these configurations. Both functional lateralization and its structural substrates change across the lifespan, and atypical organization may carry different consequences following developmental perturbation or acquired brain injury. Importantly, altered lateralization should not automatically be interpreted as adaptive or maladaptive, as the behavioral significance of hemispheric specialization varies with the affected system, developmental stage and measurement context (65). Longitudinal multimodal studies combining repeated assessment of language and attention with diffusion imaging, cortical microstructure and cognition will be necessary to establish how the phenotypes identified here emerge across development and whether individuals move between organizational states over time.

In summary, these findings support a view of hemispheric specialization as a multivariate and polymorphic property of human brain organization. Attention varied among right-dominant, bilateral and left-dominant phenotypes with distinct task-evoked and intrinsic connectivity profiles. Across domains, language and attention were related, but this relationship was expressed primarily through lateralization strength and bilaterality rather than deterministic assignment to opposite hemispheres. These profiles were heritable and accompanied by focal cortical and widespread white-matter differences, yet supported broadly comparable cognitive performance. Hemispheric specialization may therefore be better understood not as adherence to a single optimal template, but as a family of biologically constrained architectures through which distributed cognitive functions achieve specialization while preserving behavioral robustness. Further study of the relationship between cross-domain lateralization and broader features of brain organization has the potential to illuminate how cerebral dominance emerges in humans, why it varies so widely across individuals, and when that variation becomes behaviorally or clinically consequential (65, 72, 89).

## Methods

### HCP Participants

Participants were part of the April 2018 update of the S1200 release of the WU-Minn Human Connectome Project (HCP), which has been described in detail previously (45). Of the 1,206 participants included in the release, we selected individuals with complete 3 T resting-state fMRI data and complete task-fMRI data for the 2-back versus 0-back contrast of the HCP working-memory paradigm. This procedure yielded a final sample of 987 participants, including 524 females. Mean age was 28.71 years (SD=3.69 years, *n*=987), and mean educational level was 14.97 years (SD=1.76 years, *n*=986).

Manual preference was assessed using the HCP version of the Edinburgh Handedness Inventory (90). Participants with a score below 30 were classified as left-handed, whereas those with scores of 30 or higher were classified as right-handed (91, 92). Edinburgh Handedness Inventory scores are reported in the restricted HCP demographic dataset under the variable Handedness and are available only to authorized users. Because the HCP implementation includes one foot-preference item among its ten items, the resulting score does not provide an exclusive measure of manual preference (93). The final sample included 108 left-handed participants, of whom 49 were females, approximating the prevalence of left-handedness in the general population (92, 94).

HCP data collection was approved by the participating institutional review boards in the United States and Europe, under the coordination of Washington University in St. Louis and the University of Minnesota. The present study was approved by the Rutgers University Institutional Review Board.

### Cognitive Performance

Cognitive performance was assessed using unadjusted scores from the NIH Toolbox Cognition Battery. This battery includes seven individual instruments assessing language, executive function, episodic memory, processing speed, and working memory, together with Total, Crystallized, and Fluid Cognition Composite scores (95–101). Crystallized cognition primarily reflects acquired language-related abilities, whereas fluid cognition encompasses executive function, episodic memory, processing speed, and working memory.

Ten cognitive outcomes were examined: the Total Cognition Composite, Crystallized Cognition Composite, Fluid Cognition Composite, Picture Sequence Memory, Dimensional Change Card Sort, Flanker Inhibitory Control and Attention, Oral Reading Recognition, Picture Vocabulary, Pattern Comparison Processing Speed, and List Sorting Working Memory scores. These measures were respectively interpreted as indexing overall cognition, crystallized cognition, fluid cognition, visual episodic memory, cognitive flexibility, inhibitory control, reading decoding, vocabulary comprehension, processing speed, and working memory. The analysis used the corresponding unadjusted HCP variables and included 974 participants with complete cognitive, phenotype, demographic, and anatomical data.

### MRI Data Acquisition and Preprocessing

MRI data were acquired as part of the WU-Minn Human Connectome Project using multiband echo-planar imaging on a customized 3 T Siemens Skyra scanner. High-resolution structural images were acquired at 0.7 *mm* isotropic resolution. Resting-state fMRI consisted of four runs collected across the REST1 and REST2 sessions, with one left-to-right and one right-to-left phase-encoding run per session. Each run lasted 14 *min* 33 *s* and was acquired with a repetition time of 720 ms, an echo time of 33.1 *ms*, and 2 *mm* isotropic voxels. Task-fMRI data were acquired using the same multiband EPI sequence. Acquisition and preprocessing procedures for the HCP imaging protocols have been described previously (47, 102–104).

The present analyses used the *2*-back *minus 0*-back contrast from the HCP working-memory task. Relative to the lower-load *0*-back condition, the *2*-back condition increases demands on working-memory maintenance together with the sustained goal-directed attention and cognitive control required for successful performance (48, 49). We therefore used this contrast to index task-evoked attentional-control demands within the independently defined dorsal and ventral attention networks (14). Participant-level, HCP-preprocessed level-2 contrast maps were obtained in the MSMAll-aligned 32k_fs_LR cortical surface space.

For the resting-state analyses, HCP-provided cleaned volumetric data were obtained for all four runs: REST1_LR, REST1_RL, REST2_LR, and REST2_RL. Resting-state images were represented in 2 *mm* MNI volumetric space. Task and resting-state data were retrieved from the HCP Amazon Web Services repository. Thus, task-derived markers were calculated in MSMAll surface space, whereas intrinsic connectivity markers were calculated from volumetric resting-state images in MNI space.

Cortical myeloarchitecture was assessed using the HCP-provided bias-corrected and smoothed myelin maps in MSMAll-aligned fsLR32k surface space. Cortical thickness and curvature maps registered to the same surface space were also obtained for each participant and were subsequently used to account for local morphological contributions to the myelin signal.

Structural connectivity was examined using participant-level 400 by 400 connectivity matrices defined according to the homotopic cortical parcellation of Yan and colleagues (59) and its assignment to the seven network organization of Yeo and colleagues (63). We used median hindrance-modulated orientational anisotropy matrices as a tract-specific index of fiber density between cortical parcels.

### Attention Atlas Statistics

Attention-network measures were derived using the Atlas for Lateralized Attentional Networks, or ALANs (14). The atlas delineates two functionally lateralized systems: a dorsal network and a ventral network. ALANs regions were represented using the 400 parcel homotopic cortical parcellation of Yan and colleagues (59), registered separately to the fsLR32k surface and MNI volumetric spaces. Yan parcels showing substantial spatial overlap with an ALANs network, defined by an overlap coefficient greater than 0.50, were retained. The corresponding contralateral homotopic parcels were used to characterize the organization of each network in the opposite hemisphere.

Task-evoked activation was quantified from the *2*-back *minus 0*-back contrast separately for the dorsal and ventral networks. Measures were calculated both across the full network and within the principal network hubs. The dorsal network hub was located in the inferior frontal sulcus. Ventral network hubs included inferior frontal regions encompassing the pars triangularis and anterior insula, together with the posterior superior temporal sulcus. Because task activation maps were represented on the cortical surface, network activation was summarized while accounting for differences in parcel surface area. Within each selected parcel, activation was first averaged across vertices. Parcel-level activation estimates were then combined across the network using each parcel’s total surface area as a weight. For each network or hub set, hemispheric asymmetry was expressed as the difference between activation in the left and right hemispheres. Positive values therefore indicated greater left-hemisphere activation, whereas negative values indicated greater right-hemisphere activation. Four task-derived measures were retained for phenotype identification: dorsal network asymmetry, dorsal network hub asymmetry, ventral network asymmetry, and ventral network hub asymmetry.

Intrinsic functional connectivity was computed separately for each of the four resting-state runs. Within each ALANs associated parcel, a representative BOLD time series was obtained by averaging across all parcel voxels. Pearson correlation coefficients were then calculated between the parcel time series to generate one functional connectivity matrix per participant and run. The four run-specific matrices were transformed using Fisher’s z-transformation, averaged, and r-transformed back to correlation coefficients using a hyperbolic tangent function, thereby producing a single participant level connectivity matrix with improved signal-to-noise ratio and reliability (105). Three resting-state measures were derived for each attention network. First, intrahemispheric connectivity strength, or centrality degree, was calculated. For each parcel, strength was defined as the sum of correlations with the other parcels belonging to the same network and hemisphere. Parcel strengths were subsequently averaged to obtain one network-strength estimate for each hemisphere. Second, strength asymmetry was calculated as left hemisphere minus right hemisphere network strength, such that positive values represented greater leftward intrinsic connectivity and negative values represented greater rightward connectivity. Third, network strength sum was defined as the mean of the left- and right-hemisphere strength estimates. Finally, homotopic interhemispheric connectivity was quantified from correlations between corresponding parcel pairs in the two hemispheres. Homotopic correlations were Fisher z-transformed, averaged across parcel pairs, and r-transformed back.

The dorsal and ventral networks were consequently each characterized by three resting-state measures: network level strength asymmetry, network level strength sum, and network level homotopic correlation. Together with the four task-evoked asymmetry measures, these variables yielded ten functional markers describing the task-evoked and intrinsic organization of attention networks for each participant.

### Elaboration of the Cross-Domain Crowding Phenotype

Cross-domain hemispheric organization was examined in 987 participants with complete attention and language phenotype assignments. Attention phenotype was categorized as Typical, Bilateral, or Atypical. The Typical attention phenotype was characterized by rightward task-evoked asymmetry, the Atypical phenotype by leftward asymmetry, and the Bilateral phenotype by no dominant hemisphere. Language phenotype was categorized as Strong Typical, Mild Typical, or Atypical. Strong Typical and Mild Typical language phenotypes both reflected left-hemisphere language dominance, differing in the strength of their lateralization, whereas the Atypical phenotype reflected right-hemisphere language dominance (22).

A three-level crowding phenotype was derived from the joint attention and language classifications (Fig. 1-e). Participants were classified as Split when the two lateralized functions were dominant in opposite hemispheres. This category included participants with Typical attention and either Strong Typical or Mild Typical language, as well as participants with Atypical attention and Atypical language. Participants were classified as Crowded when attention and language were dominant in the same hemisphere. This category included participants with Typical attention and Atypical language, and participants with Atypical attention and either Strong Typical or Mild Typical language. Participants with Bilateral attention were classified as Mixed, irrespective of language phenotype, because no dominant attention hemisphere could be assigned.

### Statistical Analyses

Statistical analyses were performed using R (106) (R version: 4.5.0). The neuroimaging data were handled and processed using R libraries *ciftiTools* (R package version: 0.19.0) (107), *RNifti* (R package version: 1.9.0) (108), and *neurohcp* (R package version: 0.11.0) (109). Data wrangling was performed using the R library *dplyr* (R package version: 1.2.1) (110). Brain visualizations were realized using *Surf Ice* (version: 6 October 2021) (111), and were made reproducible following guidelines to generate programmatic neuroimaging visualizations (112). Other figures were made using *ggplot2* (R package version: 4.0.3) (113) and *circlize* (R package version: 0.4.16) (114).

#### Identification of attention phenotypes

Attention phenotypes were identified in 987 participants using ten participant-level measures characterizing the task-evoked and intrinsic functional organization of the dorsal and ventral attention networks. For each network, two measures were derived from the *2*-back *versus 0*-back task contrast: activation asymmetry averaged across the complete network and activation asymmetry averaged across its principal hubs. Three additional measures were derived from resting-state functional connectivity: mean intrahemispheric functional connectivity strength, asymmetry in functional connectivity strength, and homotopic interhemispheric connectivity. Asymmetry measures were calculated as the difference between the left and right hemispheres, such that positive values indicated leftward asymmetry and negative values indicated rightward asymmetry. Together, these measures provided a multivariate description of each participant’s attention-network organization across task and resting-state conditions. Before classification, each functional measure was robustly standardized by subtracting its sample median and dividing by its median absolute deviation, reducing the influence of outliers (115). Attention phenotypes were then derived using agglomerative hierarchical clustering (116). Euclidean distances were calculated between participants, and clusters were successively merged using Ward’s criterion (117). The number of clusters was determined using the R library *NbClust* (R package version: 3.0.1) (55), considering solutions from two clusters up to n0.3 (n being the number of observations) (118) and all available internal validity indices. Among indices, five supported a two-cluster solution, twelve supported a three-cluster solution, one supported four clusters, one supported six clusters, and two supported eight clusters. By majority rule, the three-cluster solution was therefore selected. Cluster identities were assigned after examining their multivariate functional profiles, particularly the direction and magnitude of task-evoked asymmetry. The cluster characterized by predominantly rightward task asymmetry was designated Typical, the cluster characterized by no dominant task-evoked hemispheric asymmetry was designated Bilateral, and the cluster characterized by predominantly leftward task asymmetry was designated Atypical.

To characterize differences in the functional profiles of the three attention phenotypes, each of the ten task-evoked and resting-state measures was subsequently analyzed using analysis of covariance, with attention phenotype entered as a categorical factor, and age, self-reported sex, manual preference, years of education, and total intracranial volume included as covariates, as well as the manual preference by attention phenotype interaction. Overall phenotype effects were evaluated using Type III analyses of variance with sum-to-zero contrasts, and p values for the phenotype effect were corrected across the ten functional measures using the Benjamini-Hochberg false discovery rate procedure. Estimated marginal means and 95% confidence intervals were obtained using the *emmeans* R library (R package version: 1.11.0) (119). Pairwise comparisons among Typical, Bilateral, and Atypical phenotypes were performed using Tukey-adjusted contrasts.

#### Association of Self-Reported Sex and Manual Preference with Attention Phenotype

To determine whether attention phenotype membership varied as a function of self-reported sex or manual preference, we fitted multinomial logistic regression models using the R package *nnet* (R package version: 7.3-20) (120). Attention phenotype, comprising Typical, Bilateral, and Atypical organization, was entered as the categorical outcome, with the Typical phenotype as the reference category. Self-reported sex and manual preference were entered as categorical predictors, with female participants and right-handers as the respective reference groups. We first fitted a model including the main effects of self-reported sex and manual preference and their interaction. Overall effects were evaluated using likelihood-ratio chi-square tests implemented from the R package *car* (R package version: 3.1-5) (121). Because the interaction was not significant, the final model included only the additive effects of self-reported sex and manual preference, which were assessed using Type II likelihood-ratio tests. Individual coefficients contrasting Bilateral and Atypical organization with the Typical reference phenotype were evaluated using two-sided Wald tests. Coefficients were exponentiated to obtain relative risk ratios (RRR), with 95% Wald confidence intervals. For visualization, model-adjusted probabilities of membership in each phenotype were calculated using the R package *marginaleffects* (R package version: 0.32.0) (122). Probabilities associated with self-reported sex were averaged over the observed distribution of manual preference, and probabilities associated with manual preference were averaged over the observed distribution of self-reported sex.

#### Statistical Assessment of Cross-Domain Organization

We evaluated cross-domain organization at two complementary levels. First, to test whether the full attention and language phenotype classifications were statistically independent, we constructed a 3×3 contingency table and performed a Pearson *χ*^2^ test of independence. Association magnitude was quantified using Cramér’s V *ϕ*_C_.

Cells contributing to the association were identified using adjusted standardized residuals, also known as Haberman residuals. Two-sided cellwise *p* values were obtained by comparing each residual with the standard normal distribution. The nine resulting *p* values were corrected for multiple comparisons using the Benjamini-Hochberg false discovery rate procedure (123). Cells with *p*_FDR_<0.05 and positive residuals were interpreted as enriched relative to the frequency expected under independence, whereas cells with *p*_FDR_<0.05 and negative residuals were interpreted as depleted.

Second, we conducted a directional test designed specifically to evaluate the complementary specialization hypothesis. Participants with Bilateral attention were excluded because they could not be assigned a dominant attention hemisphere. Strong Typical and Mild Typical language phenotypes were combined into a left-dominant language category (as previously done by Labache and colleagues (22, 23)), and Atypical language was categorized as right-dominant. The Typical and Atypical attention phenotypes were categorized as right- and left-dominant attention, respectively. A 2×2 Pearson *χ*^2^ test without continuity correction was then used to determine whether language dominance was associated with attention dominance. Opposite-hemisphere combinations represented complementary or Split organization, whereas same-hemisphere combinations represented Crowded organization. Effect magnitude was summarized using the *ϕ*_C_ coefficient and the odds ratio with its 95% confidence interval.

#### Effect of Crowding on Cognitive Performance

Analysis of covariance was used to assess the broader relationship between cross-domain crowding phenotype and cognitive performance. For each cognitive outcome, the model was specified as follows: cognitive performance was the dependent variable, cross-domain crowding phenotype, comprising Split, Mixed, and Crowded groups, was the independent variable, and age, years of education, total intracranial volume, self-reported sex, and manual preference were included as covariates. The main crowding phenotype effect therefore tested whether adjusted cognitive performance differed among Split, Mixed, and Crowded participants. Secondary analyses additionally included the interaction between manual preference and crowding phenotype to test whether the relationship between cross-domain organization and cognitive performance differed according to handedness. Sum-to-zero contrasts were used for categorical predictors, and effects were evaluated using Type III analyses of variance. Effect sizes were quantified using partial eta-squared (*η*_p_ ^2^). The Benjamini-Hochberg false discovery rate procedu^p^re (FDR) (123) was used to correct for the 10 cognitive outcomes, separately for the crowding-phenotype main effects and crowding phenotype by manual preference interactions. For outcomes showing a significant FDR corrected main effect, post hoc comparisons between crowding phenotypes were conducted using Tukey’s range test.

#### Heritability of Attention and Crowding Phenotype

Heritability of attention and cross-domain phenotype membership was estimated separately using the multidimensional estimator described by Anderson and colleagues (57). Analyses included 983 HCP participants from 444 families, comprising 129 monozygotic twin pairs (*n*=258), 69 dizygotic twin pairs (*n*=138), 475 non-twin siblings and 112 unrelated singletons. For each phenotype, we constructed a participant by participant similarity matrix with entries of 1 for participants assigned to the same category and 0 otherwise. The additive genetic relationship matrix, containing twice the pedigree kinship coefficients, was calculated using SOLAR (124) through the R package *solarius* (125). Analyses adjusted for age, sex, age2, age by sex, age^2^ by sex, handedness and intracranial volume. Age, age^2^ and intracranial volume were standardized before model fitting. Covariate adjustment was performed by projecting both similarity and relationship matrices into the subspace orthogonal to the covariate design matrix, which included an intercept. Heritability was estimated by moment matching as the ratio of the traces of the estimated additive genetic and total phenotypic covariance matrices, with estimates truncated to the interval [0, 1]. Unadjusted, one-sided permutation p-values were calculated using 999 random simultaneous permutations of the rows and columns of the projected relationship matrix, together with the observed arrangement, yielding 1,000 reference estimates. *P* values were calculated as the proportion of reference estimates greater than or equal to the observed heritability. Standard errors were estimated using a leave-one-family-out block jackknife, removing all participants belonging to each family in turn and recalculating heritability. The model included additive genetic and residual components without a separate shared environmental component.

#### Effect of Crowding on Myelination

Bias-corrected HCP cortical myelin maps, cortical thickness maps, and curvature maps were analyzed in the fsLR32k surface space. Bilateral surface data were assembled into participant-by-vertex matrices using the R library *VertexWiseR* (R package version: 1.4.1) (126). *VertexWiseR* supports direct extraction and analysis of fsLR32k data generated by the HCP processing pipeline. Participants with missing demographic or phenotype data were excluded, resulting in a final sample of 986 participants: 437 Split, 385 Mixed, and 164 Crowded.

To reduce potential contributions of cortical morphology to the myelin signal, myelin values were residualized separately at each cortical vertex (127). For each vertex, an ordinary least-squares model was fitted with the myelinmap value as the dependent variable and local cortical thickness and curvature as predictors. The residuals from these models were retained as the vertex-wise measure of cortical myelination. No additional surface smoothing was applied to these residualized data.

Whole-cortex vertex-wise linear models were fitted using the Vertex-wise analysis with random field theory (RFT) cluster correction function (RFT_vertex_analysis) in the *VertexWiseR* library. This function fits linear models through BrainStat and applies random field theory-based cluster correction. The model included age, self-reported sex, total intracranial volume, manual preference, years of education, and crowding phenotype. Identified significant clusters were assessed separately at a cluster-level RFT-corrected threshold of *p*<0.05. *VertexWiseR* reports the cluster extent, cluster-corrected probability, peak *t* statistic, and MNI coordinates of each significant cluster.

For visualization and characterization of the significant cluster, each participant’s mean residualized myelin value was calculated across vertices belonging to the significant cluster mask. These values were analyzed using a linear model that included crowding phenotype as a categorical factor and age, self-reported sex, total intracranial volume, and manual preference as covariates. The overall association with crowding phenotype was evaluated using a Type III analysis of variance with sum-to-zero contrasts. Estimated marginal means were obtained using the emmeans, and all pairwise comparisons among Split, Mixed, and Crowded phenotypes were adjusted using Tukey’s range test.

#### Crowding Effect on Structural Connectivity

Participant-level structural connectivity matrices were analyzed using a cortical parcellation comprising 400 homotopic parcels assigned to the seven-network organization proposed by Yeo and colleagues (59, 63). The pre-processed diffusion MRI data for each participant were modelled in StarTrack (https://nbl-research.github.io/) using the damped Richardson-Lucy spherical deconvolution algorithm (62, 128) applied to the top two shells (*i.e., b*=2,000 and *b*=3,000 *s*/*mm*^2^) according to the following parameters: fiber response *α*=1.5; number of iterations=200; amplitude threshold *η*=0.0015; geometric regularisation *ν*=16. Fibre tracking was then performed using the multi-fibre Euler-like algorithm with the following parameters: minimum hindrance-modulated orientational anisotropy (HMOA) threshold=0.003; step size=1 *mm*; maximum angle threshold=45; minimum fibre length=20 *mm*; maximum fibre length=300 *mm*. The resulting whole-brain tractogram was converted to a connectivity matrix by projecting the streamline terminations to the nearest vertex on the cortical surface within 2.5 *mm*, and assigning the median HMOA value of all streamlines connecting a pair of parcels to the corresponding edge. Therefore, each matrix was symmetric and contained the median HMOA value for each parcel-to-parcel connection. HMOA was interpreted as a tract-specific index of fiber density. Participants were matched across the structural connectivity, demographic, attention phenotype, and language phenotype datasets using their HCP participant identifiers. Participants without complete demographic or phenotype information were excluded. The final structural-connectivity sample comprised 981 participants: 433 Split, 384 Mixed, and 164 Crowded.

Because structural connectivity matrices were symmetric, statistical analyses were restricted to the upper triangular portion of each matrix. Connections for which HMOA was zero in every participant were excluded, leaving 62,182 nonzero edges for analysis. For each edge, an ordinary least-squares linear model was fitted with HMOA as the dependent variable. HMOA was the independent variable, and crowding phenotype, age, years of education, total intracranial volume, self-reported sex, and manual preference were included as covariates, as well as the manual preference by crowding phenotype interaction.

Categorical predictors were represented using sum-to-zero contrasts, and the main effect of crowding phenotype was evaluated using a Type III analysis of variance. The resulting p values were corrected across all nonzero edges using the Benjamini-Hochberg false discovery rate procedure. Connections with *p*_FDR_<0.05 were considered significant.

Significant edges were annotated according to the seven-network assignments of their constituent parcels. Network pairs were treated as undirected, such that, for example, dorsal-attention-to-default and default-to-dorsal-attention connections belonged to the same category. To characterize the direction and magnitude of the crowding-phenotype effect, HMOA was averaged for each participant across all significant edges belonging to a given network pair. These participant-level averages were analyzed using the same linear-model specification as the edge-wise analyses. Estimated marginal means for the Split, Mixed, and Crowded phenotypes were obtained, and pairwise comparisons were adjusted using Tukey’s range test within each network pair.

To account for differences in network size, the number of significant edges within each network pair was normalized by the number of anatomically possible connections. For two different networks containing *n*_A_ and *n*_B_ parcels, the number of possible connections was *n*_A_×*n*_B_.

For a within-network comparison involving *n*_A_ parcels, the number of possible connections was ((*n*_A_(*n*_A_ *-1))/2)*. Network-level involvement was summarized by combining the normalized edge densities of all network pairs incident on each network, with within-network values counted once.

## Data Availability Statement

The HCP data analyzed in this study are available through the Human Connectome Project under its applicable access terms. The analyzed participant list, derived nonrestricted data and figure source data are available at: https://github.com/loiclabache/Labache_2026_HeCo. Restricted demographic and family information must be obtained through the HCP access process.

## Code Availability Statement

The code used in the Method section to reproduce the results, and the visualizations can be found here: https://github.com/loiclabache/Labache_2026_HeCo.

## Acknowledgments

This work was supported by the National Institute of Mental Health (*R01MH120080*, and *P50MH136296* to AJH), and by Rutgers Health Center for Biomedical Informatics and Health AI (*BMIHAI 2025 Postdoctoral Fellowship* to LL).

## CRediT Authorship Contribution Statement

**Loïc Labache:** Conceptualization, Data Curation, Formal Analysis, Investigation, Methodology, Resources, Software, Validation, Visualization, Writing—original draft, Writing—review & editing, Supervision, and Project administration. **Ahmad Beyh:** Data Curation, Investigation, Methodology, Resources, Software, Validation, Visualization, Writing—review & editing. **Rachel Cooper:** Methodology, Software, Visualization, Validation, Writing—review & editing. **Linden Parkes:** Validation, Visualization, Writing—review & editing. **Avram J. Holmes:** Conceptualization, Funding acquisition, Investigation, Methodology, Project administration, Resources, Supervision, Validation, Visualization, Writing—original draft, Writing—review & editing.

## Competing Interests Statement

The authors declare no actual or potential conflict of interest.

## Notes

https://github.com/loiclabache/Labache_2026_HeCo

### Competing Interest Statement

The authors have declared no competing interest.

